# TMS alters multivoxel patterns in the absence of overall activity changes

**DOI:** 10.1101/2020.03.25.008334

**Authors:** Farshad Rafiei, Martin Safrin, Martijn E. Wokke, Hakwan Lau, Dobromir Rahnev

## Abstract

Transcranial magnetic stimulation (TMS) has become one of the major tools for establishing the causal role of specific brain regions in perceptual, motor, and cognitive processes. Nevertheless, a persistent limitation of the technique is the lack of clarity regarding its precise effects on neural activity. Here, we examined the effects of TMS intensity and frequency on concurrently recorded blood-oxygen level-dependent (BOLD) signals at the site of stimulation. In two experiments, we delivered TMS to the dorsolateral prefrontal cortex in human subjects of both sexes. In Experiment 1, we delivered a series of pulses at high (100% of motor threshold) or low (50% of motor threshold) intensity, whereas in Experiment 2, we always used high intensity but delivered stimulation at four different frequencies (5, 8.33, 12.5, and 25 Hz). We found that the TMS intensity and frequency could be reliably decoded using multivariate analysis techniques even though TMS had no effect on overall BOLD activity at the site of stimulation in either experiment. These results provide important insight into the mechanisms through which TMS influences neural activity.

**Significance:** Transcranial magnetic stimulation (TMS) is a promising tool for the treatment of a number of neuropsychiatric disorders. However, its effectiveness is still impeded by an incomplete understanding of its neural effects. One fundamental unresolved issue is whether TMS leads to local changes in overall neural activity in the absence of a task. Here we performed two experiments where TMS was delivered inside an MRI scanner while brain activity was continuously monitored. We found converging evidence for the notion that TMS affects the pattern of local activity changes but does not lead to an overall increase in activity. These results help clarify the mechanisms of how TMS affects local neural activity.

## Introduction

Transcranial magnetic stimulation (TMS) is one of the major tools commonly used in both clinical interventions and basic research on the functions of different brain areas (Pasley et al., 2009; Li et al., 2017; Romero et al., 2019a). However, although much progress has been made, the precise effects of TMS on neural activity remain poorly understood thus limiting progress in clinical as well as basic research applications. One promising technique that can reveal both the local and distant effects of TMS is the application of TMS inside an MRI scanner while concurrently collecting functional MRI (fMRI) data (for reviews, see Bestmann and Feredoes, 2013; Bergmann et al., 2016; Polanía et al., 2018).

The feasibility of combining TMS with fMRI was first shown more than 20 years ago (Bohning et al., 1997) and the technique of concurrent TMS-fMRI has further gained in popularity since then (Bestmann and Feredoes, 2013; Heinen et al., 2014a; Leitão et al., 2015; Hawco et al., 2018; Vink et al., 2018a). Most of the initial studies using concurrent TMS-fMRI examined the effects of TMS delivered to the primary motor cortex (M1). This choice was partially driven by the fact that localization of M1 is straightforward and can be ascertained by observing contralateral hand muscle twitches. One ubiquitous finding from these studies is that TMS induces increased blood-oxygen level-dependent (BOLD) activity in the vicinity of the targeted area for suprathreshold (Bohning et al., 1999; Baudewig et al., 2001; Bestmann et al., 2004) but not subthreshold stimulation intensities (Bohning et al., 2000b; Baudewig et al., 2001; Bestmann et al., 2003, 2004, 2005; Li et al., 2004b; Denslow et al., 2005b; Navarro de Lara et al., 2017). These findings led researchers to conclude that the activation produced by suprathreshold intensities is likely the result of afferent feedback arising from the induced twitches (Baudewig et al., 2001; Bestmann et al., 2003, 2006; Bestmann and Feredoes, 2013). Similar results have been found when delivering TMS over visual cortex: local activity increase is observed only for intensities high enough to produce phosphene perception (Caparelli et al., 2010), thus suggesting that the observed BOLD activity might be the result of feedback from higher cortical regions responsible for the conscious experience of the phosphenes. These studies highlight the difficulties in characterizing local TMS effects on neural activity independent of their downstream consequences (e.g., twitches or phosphene perception).

Therefore, understanding the local effects of TMS independent of overt behavioral responses such as twitches and phosphenes requires that TMS is applied outside of the motor and visual cortices. Several studies have targeted areas in the association cortex but were typically not designed specifically to investigate whether TMS produces activations in the targeted area. Consequently, the results of these studies were inconsistent: some studies reported that in the absence of an explicit task, TMS produces positive local BOLD activations (Nahas et al., 2001; Li et al., 2004a; Bestmann et al., 2005; Vink et al., 2018b) but many other studies found no such activations (Baudewig et al., 2001; Kemna and Gembris, 2003; Li et al., 2004b; Sack et al., 2007; De Weijer et al., 2014; Hawco et al., 2017; Dowdle et al., 2018). These previous studies generally conducted whole-brain general linear model (GLM) analyses. Such analyses are most powerful if the stimulation location is perfectly matched across subjects, but this is known not to be the case in concurrent TMS-fMRI studies where the true location of the TMS coil typically deviates somewhat from the desired one (Leitão et al., 2017). This induces variability in stimulation location between subjects and can increase the chance of observing both false positives and false negatives in standard GLM analyses. Therefore, to convincingly establish the local effects of stimulation, a more powerful approach would be to identify the exact location of stimulation for each subject as a region of interest (ROI) and then perform all analyses on these ROIs.

Here we investigated the local effect of TMS applied to the dorsolateral prefrontal cortex (DLPFC). We adopted the above approach by precisely localizing the stimulated site as an ROI for each subject and performing the main analyses on these ROIs. We also delivered a large number of pulses for each participant (1,200 pulses per subject in Experiment 1 and 1,800 pulses per subject for each of the two days of stimulation in Experiment 2) to ensure that we can obtain meaningful results on an individual-subject level. DLPFC was chosen because it is easily accessible inside the MRI scanner and stimulating it does not result in any motor or visual responses. In Experiment 1, we compared the activations produced by low-intensity 1 Hz trains (10 pulses each), high-intensity 1 Hz trains (10 pulses each), and high-intensity bursts of 4 pulses delivered at 0.5 Hz (20 pulses each). In Experiment 2, we compared the activations produced by trains of 30 pulses delivered at four different frequencies (5, 8.33, 12.5, and 25 Hz). Despite the large number of overall pulses delivered in both experiments, we found no evidence that TMS increased local BOLD activity. However, we were able to decode both the intensity (Experiment 1) and frequency (Experiment 2) of stimulation using multivoxel pattern analysis (MVPA). These results demonstrate that different TMS stimulation protocols produce different patterns of activity in the underlying areas in the absence of an overall increase in BOLD activity.

## Methods

### Subjects

Five subjects (2 females, age range = 24-33) took part in Experiment 1 and six subjects (3 females, age range = 24-37) took part in Experiment 2. Three subjects participated in both of the experiments. Unlike many cognitive studies that seek to aggregate data across subjects, here we were interested in the effects at the level of each individual subject. Therefore, as is common in this type of research (e.g., Rahnev et al., 2013; Huth et al., 2016), we collected data from fewer subjects but tested each of them extensively. All subjects were medically and neurologically healthy with no history of brain injury, loss of consciousness, or psychiatric diseases. They were all screened for MRI and TMS exclusion criteria and signed an informed consent document. All procedures were approved by the Georgia Tech Institutional Review Board.

### Procedures

#### Experiment 1

In order to determine the local effects of TMS, we employed three different TMS conditions (**Figure 1A**). The first condition consisted of 10 low-intensity (50% of motor threshold) TMS pulses given at a rate of 1 Hz. The second condition consisted of 10 high-intensity (100% of motor threshold) TMS pulses given at a rate of 1 Hz. Finally, the third condition consisted of five bursts of four high-intensity (100% of motor threshold) TMS pulses with individual pulses in a single burst delivered at 12.5 Hz (i.e., there was 80 ms between two consecutive pulses) and individual bursts delivered at 0.5 Hz (i.e., consecutive bursts started 2 seconds apart). Each condition was created to last 10 seconds. We call the 10-second period of TMS presentation a “trial.” Each trial thus consisted of 10 TMS pulses for Conditions 1 and 2, and 20 TMS pulses for Condition 3.

**Figure 1.**
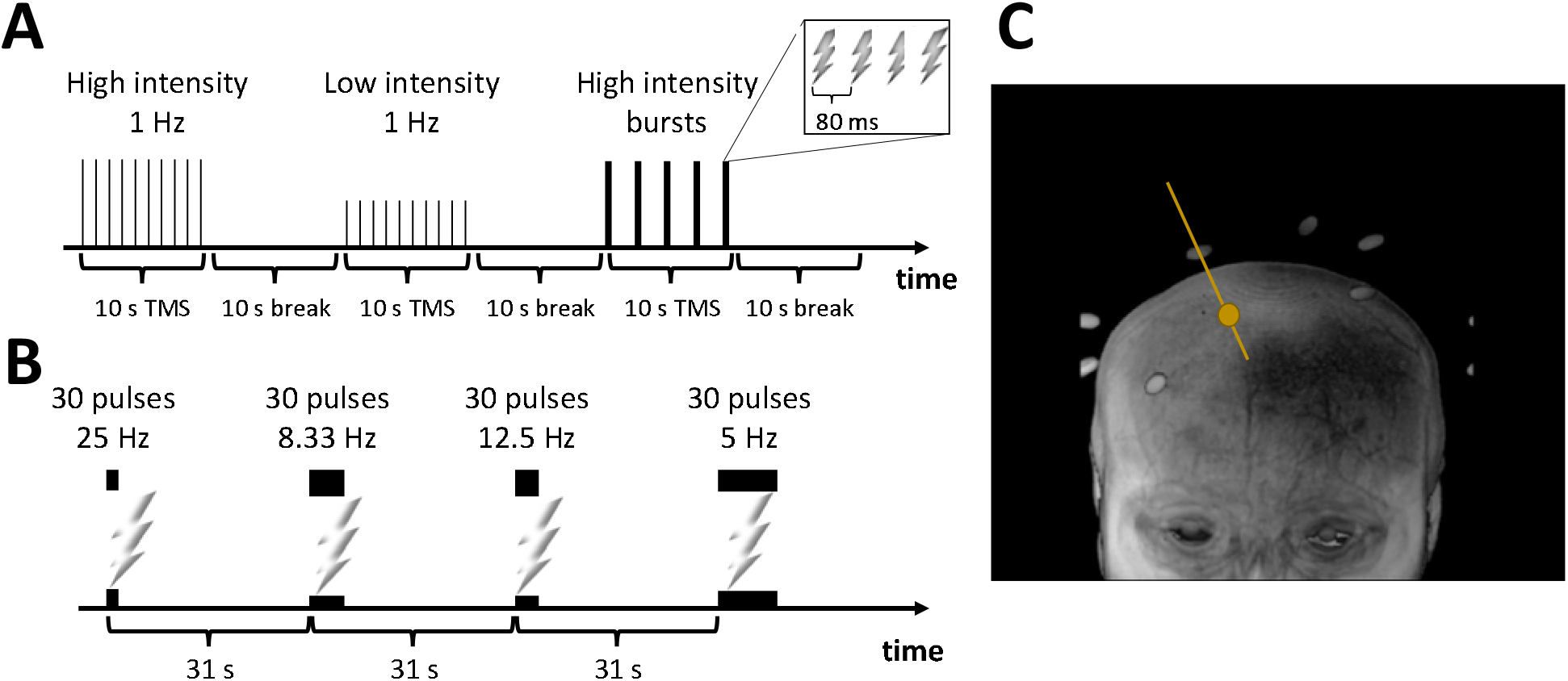
TMS delivery in Experiments 1 and 2. (A) An example block from Experiment 1. In each block, three conditions were interleaved: Low intensity 1 Hz, High intensity 1 Hz, and High intensity bursts. Each burst in the High intensity bursts condition consisted of 4 pulses with 80 ms between pulses (see inset). Both 1 Hz conditions delivered 10 pulses with 1 second between consecutive pulses, whereas the High intensity bursts condition consisted of 5 bursts (with 2 seconds between the onsets of consecutive bursts). The vertical lines indicate individual TMS pulses and their height indicates intensity. The thicker lines in the High intensity bursts condition indicate the presence of multiple TMS pulses in close proximity. Each TMS presentation lasted 10 seconds and a block lasted 60 seconds. (B) Four conditions were interleaved in Experiment 2: 5, 8.33, 12.5, and 25 Hz stimulation. One trial from each condition consisted of 30 pulses and the onset asynchrony between consecutive trials was 31 seconds. The black boxes indicate the period of stimulation for each trial. (C) Estimation of the stimulation spot. The image shows the anatomical image for one subject. The bright points outside of the brain are the Vitamin E capsules attached to the TMS coil. The orange line is the estimated direction of stimulation computed based on the locations of the Vitamin E capsules. The orange circle indicates the stimulation spot located on the brain.

Subjects completed three runs consisting of 10 blocks each. One block lasted 60 seconds and consisted of three TMS trials (one for each of the three conditions) with 10-second breaks after every trial (**Figure 1A**). Overall, each run consisted of 30 trials and the whole stimulation period lasted for 600 seconds (10 blocks × 60 seconds per block). We acquired additional 10 seconds of scanning at the beginning and 5 seconds of scanning at the end of each run, making each TMS block last a total of 615 seconds. Each run involved the delivery of 400 TMS pulses for a total of 1,200 TMS pulses over the course of the experiment. Throughout the experiment, subjects were asked to fixate on a white fixation cross presented on a gray screen.

In addition to the TMS scans, we collected two anatomical images, a finger tapping run, and a resting state run. The anatomical images were acquired in the beginning and in the end of the session to ensure that subject movement did not result in substantial change in the TMS coil position from the beginning to the end of the experiment. The data from the resting state scan were not analyzed for the purposes of the current paper. The finger tapping run was intended as a control task that can be used to evaluate the power of our setup to detect brain activations, as done in previous concurrent TMS-fMRI studies (Bestmann et al., 2003, 2005; Navarro de Lara et al., 2017; Vink et al., 2018b). During that run, subjects were asked to tap with their right index finger once every second for 10 seconds, followed by a 10-second break with no finger movement. The cue for finger tapping was the fixation cross changing color from black to white. The finger-tapping task thus had the exact same parameters as Conditions 1 and 2 of the TMS run. The finger-tapping run consisted of 15 alterations of finger tapping and rest (as well as a 10-second additional rest at the end of the run) for a total of 310 seconds.

#### Experiment 2

Experiment 2 included two testing sessions collected on different days and separated by less than one week. We interleaved four different conditions that differed in the frequency of stimulation. In each condition, 30 TMS pulses were given at a rate of 5, 8.33, 12.5, or 25 Hz, respectively (**Figure 1B**). These frequencies were chosen based on the time it takes to acquire a single fMRI slice, which in our protocol was 40 ms (see below). We delivered each TMS pulse so that a single slice would be affected. The four different conditions delivered TMS pulses every 1, 2, 3, or 5 slices thus resulting in frequencies of 25, 12.5, 8.33, and 5 Hz, respectively. The TMS intensity was set to 100% of motor threshold for all conditions. Each of the two sessions consisted of three TMS runs. A run was organized in 5 blocks, each consisting of four trials (one from each condition in a randomized order). Therefore, a total of 1,800 TMS pulses (3 runs × 5 blocks × 4 trials/block × 30 pulses per trial) were delivered in each session. We set the distance between the onsets of consecutive trials to 31 seconds (that is, exactly 25 fMRI volumes). Each run thus lasted 682 seconds. Each session consisted of anatomical image acquisition (both in the beginning and end of each session) and three TMS runs.

### TMS delivery and timing

Our TMS protocols were within the safety limits (Rossi et al., 2009). To further ensure subject safety, they were given a “TMS stopper” device with which they could immediately terminate the stimulation at any point during the experiment. No subject reported any atypical discomfort or symptoms either during or after the experiment. TMS was delivered with a magnetic stimulator (MagPro R100, MagVenture), using an MRI-compatible figure of eight coil (MRI-B90). We determined the resting motor threshold (RMT), immediately prior to starting the main experiment as in our previous work (Rahnev et al., 2016; Shekhar and Rahnev, 2018; Bang et al., 2019). We first localized the motor cortex by applying supra-threshold single pulses and determined the location of the motor cortex as the region that induced maximal finger twitches in the contralateral hand. The resting motor threshold (RMT) was then identified as the intensity which leads to contralateral muscle twitches on 5 out of 10 trials. The average RMT was 59.20 (SD = 10.06) in Experiment 1 and 60.67 (SD = 2.42) in Experiment 2.

The delivery of a TMS pulse creates a large artifact in the fMRI slice that is collected during the TMS delivery. Therefore, in Experiment 1, we delivered the TMS pulses during slices that would have as minimal as possible of an effect on later analyses. Specifically, in Conditions 1 and 2, which consisted of trains of single pulses at 1 Hz, we delivered the TMS pulse during the acquisition of the lowest brain slice (25^th^ slice in our descending sequence). Similarly, in Condition 3, which consisted of burst of 4 pulses at 80 Hz spaced out by 2 seconds, we delivered the TMS pulses when acquiring 19^th^, 21^st^, 23^rd^ and 25^th^ slices of the brain. These deep slices were never used in the current analyses, so the artifacts created there did not affect our results in Experiment 1.

However, such a scheme was not possible in Experiment 2 where 30 pulses were delivered in close temporal succession. Instead, to facilitate data cleanup, in Experiment 2 we ensured that a particular slice was never targeted in two consecutive fMRI volumes. This was achieved by making each fMRI volume with 31 slices and starting the sequence in all conditions from slice 1. Thus the 25 Hz condition targeted slices 1, 2, …, 30 from a single volume, the 12.5 Hz condition targeted slices 1, 3, 5, …, 31 in the first volume and slices 2, 4, 6, …, 28 in the second volume, etc. Therefore, the 25 Hz condition corrupted most slices in a single volume, the 12.5 Hz condition corrupted every other slice in two consecutive volumes, the 8.33 Hz condition corrupted every third slice in three consecutive volumes, and the 5 Hz condition corrupted every fifth slice in five consecutive volumes. This pattern of stimulation allowed us to clean the contaminated data with minimal loss of signal (see below).

### TMS targeting

In Experiment 1, we targeted right DLPFC by placing the TMS coil over the right dorsal frontal region of an individuals’ brain. In Experiment 2, we stimulated a site in left DLPFC that is sometimes targeted in treatments for depression (Fox et al., 2012). Specifically, the stimulation spot in Experiment 2 was defined as the voxel in the left middle frontal gyrus with largest negative connectivity with the subgenual nucleus based on a previous fMRI scan. We marked the stimulation spot in subjects’ anatomical space and used the titanium-based, MRI-compatible Localite TMS Navigator and camera stand to position the coil over the identified location. In both experiments, we ensured that our TMS site did not result in any muscle twitches by first delivering several TMS pulses immediately after positioning the coil and closely monitoring the subject’s contralateral part of the body.

Regardless of the targeting method used, we wanted to independently verify the exact location of stimulation in both experiments. Therefore, we designed a procedure where we attached seven Vitamin E capsules on the surface of the TMS coil in order to help us determine the exact placement of the coil (**Figure 1C**). Six of the Vitamin E capsules were placed on the bottom surface of the coil in a hexagon shape centered on the middle of the TMS coil. Averaging the location of all six Vitamin E capsules gave us the center of the hexagon, which allows us to determine the exact location of the center of the TMS coil. To determine the exact axis of stimulation, we used the last Vitamin E capsule, which was attached at the center of the coil but on its upper surface. The axis of stimulation (or entry line) was then computed by finding the equation of a line which passes through the seventh Vitamin E capsule and is perpendicular to the plane created by the first six capsules.

The procedure additionally allowed us to estimate how much the stimulation spot shifted over the course of each experiment. To do so, we coregistered the anatomical images collected at the beginning and end of each session and determined the stimulation spot separately for each of them. Finally, we calculated the coil displacement from the beginning to the end of the experiment as the distance between these two points.

### MRI data acquisition

Data collection was performed on a Siemens 3T Trio scanner. Since the TMS coil is too large to fit into standard MRI receiver coils, we employed a setup that consists of the bottom part of a 12-channel MRI coil together with a 4-channel FLEX coil that wraps on the top of the subject allowing us to obtain full brain coverage. High resolution T1-weighted anatomical images were acquired with MPRAGE pulse sequence (FoV = 256 mm; TR = 2250 ms; TE = 3.98 ms; 176 slices; flip angle = 9°; voxel size = 1.0 × 1.0 × 1.0 mm^3^). Functional images were acquired using a T2*-weighted gradient echo-planar imaging sequence. To increase our signal near the site of stimulation in Experiment 1, we only had partial brain coverage near the top of head and used a TR of 1,000 ms and 25 descending slices (40 ms per slice). On the other hand, Experiment 2 aimed for a full-brain coverage and used TR of 1,240 ms and 31 descending slices (40 ms per slice). All other parameters were equivalent between the two experiments (FoV = 220 mm; TE = 30 ms; flip angle = 50°; voxel size = 3.0 × 3.0 × 3.5 mm^3^).

### MRI preprocessing

All analysis steps were performed in SPM12. We first transformed the original DICOM images to NIFTI format and removed all TMS-related artifacts in the following manner. In Experiment 1, since the deepest slice of the brain (25^th^ slice) was contaminated in all stimulation conditions, we removed this slice from further analysis for all subjects. In addition, for Condition 3 of Experiment 1 and for all conditions of Experiment 2, we removed each targeted slice and substituted it with the first order interpolation (i.e., mean) of the same slice number from the previous and next volumes. Finally, to ensure that the data were free from any additional signal loss, we checked for artifacts by examining the average signal amplitude in each slice over the time course of each run of both experiments. Deviations of more than four standard deviations were flagged as outliers (less than 0.05% in Experiment 1 and less than 0.5% in Experiment 2) and removed by a first order interpolation of the same slice number from the previous and next volumes. If any of the slices used in the interpolation were also flagged as outliers, we removed both slices and used a first order interpolation of the closest slices not flagged as outliers.

After removing the TMS-related artifacts, we implemented standard preprocessing. We first performed additional despiking using the 3dDespike function in AFNI. The fMRI images were then slice-time corrected, realigned to the first volume of the run, coregistered to the anatomical image acquired at the beginning of the session (reference image), normalized to MNI anatomical standard space, and spatially smoothed with a 6 mm full width half maximum (FWHM) Gaussian kernel. The same preprocessing steps, except for the TMS artifact removal, were also performed for the finger tapping run in Experiment 1. This procedure resulted in good alignment between the anatomical and functional scans (**Supplementary Figure 1**).

### GLM analyses

In both experiments, we constructed a general linear model (GLM). In Experiment 1, the GLM consisted of 10 regressors for each run. The first three regressors modeled the BOLD responses related to each TMS trial for Condition 1 (Low intensity single pulse), Condition 2 (High intensity single pulse), and Condition 3 (High intensity burst of pulses). The regressor for the TMS trials modeled the whole 10-second period of stimulation rather than each individual TMS pulse separately. In Experiment 2, the GLM consisted of 11 regressors for each run. The first four regressors modeled the BOLD responses related to each TMS trial of the four TMS conditions with regressors again modeling the whole period of stimulation rather than individual TMS pulses. We included six regressors related to head movement (three translation and three rotation regressors) and a constant term. In all cases, the BOLD response was modeled with canonical hemodynamic response function (HRF). The activation maps for each subject were obtained using t-tests.

To further explore the effect of TMS specifically on the targeted area, we defined spherical regions of interest (ROIs) at the location of stimulation with radii of 8, 12, 16, and 20 mm using the toolbox MarsBaR (Brett et al., 2002). The centers of these spheres were placed on the TMS entry line and were set so that the entire ROI fell inside the brain. In control analyses, we defined spherical ROIs in auditory and visual cortex using the following Talairach coordinates: left auditory cortex: [−49, −20, 9], right auditory cortex: [48, −21, 10], left visual cortex: [−11, −78, 10], right visual cortex: [11, −75, 11] (Papademetris et al., 2006). We defined spherical ROIs with radii of 20 mm centered around these coordinates. We then performed inverse normalization to bring the ROIs from standard space to each subjects’ anatomical space. The same GLMs defined above were used to estimate the beta values for each condition in every ROI in both experiments.

### MVPA

In addition to the GLM analyses, we performed multi-voxel pattern analyses (MVPA) within all ROIs. To do so, we first defined an additional GLM where each TMS trial was modeled separately. In Experiment 1, we had a total of 90 trials total (3 conditions × 30 events per condition). We used Gaussian Naïve Bayes (GNB) classification with leave-one-run-out (3-fold) scheme implemented by The Decoding Toolbox (TDT) (Hebart et al., 2015). For each classification fold, beta estimates for each condition for two runs were used as training data, and the performance was tested on data from the third (‘left-out’) run. This was performed iteratively until all runs had been tested (i.e., 3 iterations total). The accuracy of the classifier was computed separately for each ROI and each subject by averaging the classification accuracy over all folds. We followed a similar procedure for the analyses in Experiment 2 and applied a GNB classifier to each session separately. We used TDT toolbox to generate the feature vectors and performed the analysis in the Waikato Environment for Knowledge Analysis (WEKA) software (Frank et al., 2016). We further examined whether a classifier trained on one pair of frequencies can be used to classify a different pair of frequencies. Specifically, we trained a classifier to distinguish between 5 and 25 Hz TMS using the voxels in 20-mm ROI defined at the location of stimulation and examined the ability of this classifier to distinguish between 8.33 and 12.5 Hz TMS.

In a separate analysis we explored whether the voxels that were most informative for classification were located near the surface of the brain. We performed dimensionality reduction on the data in the 20-mm ROI defined at the location of stimulation in order to identify the most informative voxels (i.e., the voxels with the most information with respect to classes; average number of selected voxels = 83.75 voxels, SD = 8.76 voxels). We employed information gain dimensionality reduction based on the Gini impurity index (Langs et al., 2011). Given the rapid decrease of the magnetic field away from the TMS coil, we hypothesized that the informative voxels should come preferentially from the upper hemisphere of the ROI. One potential concern with this analysis is that the upper hemisphere may contain more gray matter voxels, thus biasing the results. To check whether this was indeed the case, we determined the number of gray matter voxels in the upper and lower hemispheres and ensured that these numbers did not differ.

Statistical analyses were performed by fitting mixed models using the Nonlinear Mixed-Effects Models (*nlme*) library (Pinheiro et al., 2019) in R. The models were fit to the accuracy of the classification for each trial or to the proportion of gray matter voxels in the upper hemisphere of the 20-mm ROI defined at the location of stimulation for each subject. In the analyses for Experiment 1, subject was defined as a random effect. In the analyses for Experiment 2, both subject and session were defined as random effects. In all cases, the intercept was compared to chance level.

### Data and code

Processed data including beta values and classification for each subject and each ROI, as well as codes for analyzing these data are freely available at https://osf.io/z5jtg/.

## Results

We investigated the effect of TMS on local BOLD activity when targeting DLPFC. In Experiment 1, we employed three types of TMS protocols that differed in both intensity (50% vs. 100% of motor threshold) and frequency of stimulation (**Figure 1A**). In Experiment 2, we only used high intensity (100% of motor threshold) pulses but systematically varied the stimulation frequency (5, 8.33, 12.5, and 25 Hz; **Figure 1B**). To understand the effects of TMS, we performed conventional univariate analyses, as well as multivariate decoding.

### Experiment 1

#### Setup validation

We employed a novel concurrent TMS-fMRI setup in which MRI images were collected using a combination of the bottom half of a standard 12-channel MRI receiver coil together with a 4-channel FLEX coil positioned over subjects’ forehead. Therefore, we wanted to confirm that our new setup can detect changes in the BOLD signal in a task designed to have a similar structure as our TMS protocol. To do so, we employed a finger tapping task that has previously been used in concurrent TMS-fMRI studies as a point of reference for the effects of TMS applied to M1 (Boecker et al., 1994; Baudewig et al., 2001; Bestmann et al., 2003; Dechent and Frahm, 2003). Subjects pressed a button using their right index finger whenever a visual cue was presented on the screen with the presentation of the visual cue being identical to the 1 Hz TMS stimulation conditions such that 10 cues presented at 1 Hz alternated with 10-second blocks of rest.

Despite the fact that the finger tapping task was substantially shorter than the TMS task (5 minutes vs. 30 minutes total), we found that, compared to periods of rest, periods of finger tapping produced significant BOLD signal increase (at *p* < .05 FWE) in left M1 and supplementary motor area (SMA) for all subjects (**Figure 2**). These activations are in line with previously reported activity patterns evoked by the finger tapping task (Boecker et al., 1994; Baudewig et al., 2001; Bestmann et al., 2003; Dechent and Frahm, 2003) and demonstrate that our novel setup allows us to robustly detect cortical activations in dorsal regions of the brain.

**Figure 2.**
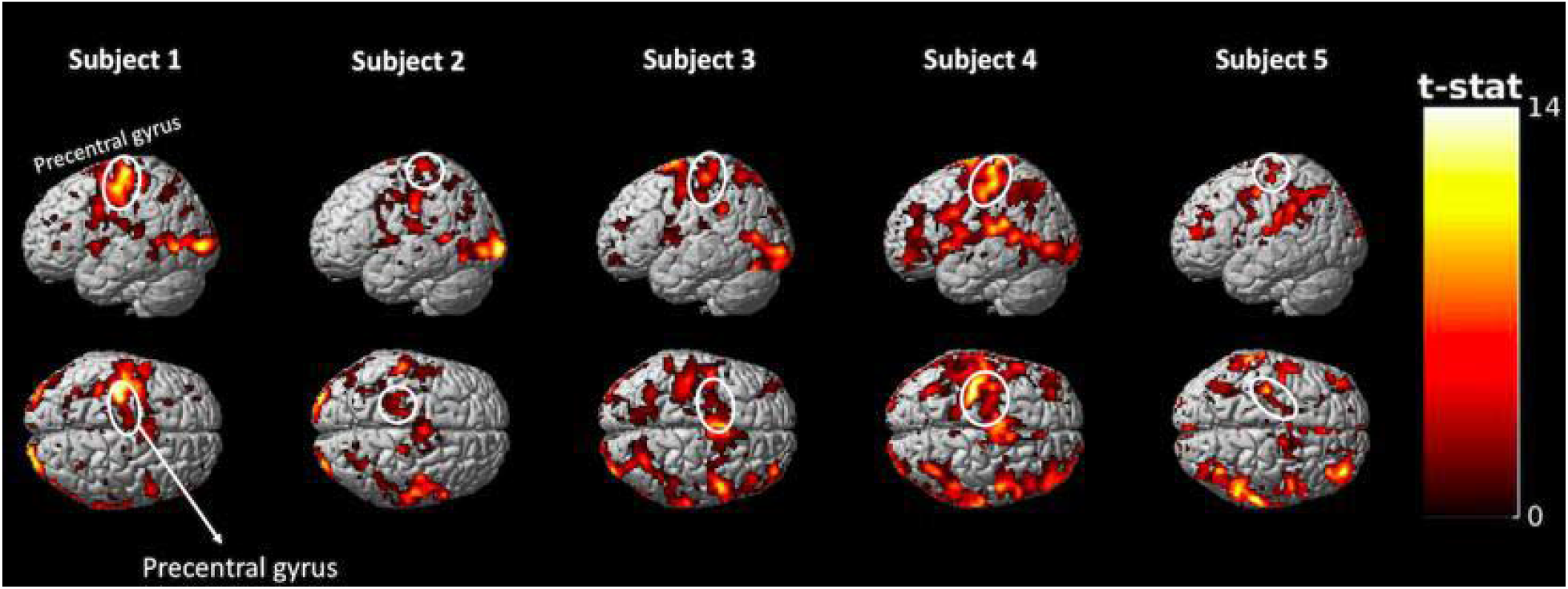
Activation maps obtained from a 5-minute finger tapping task. The finger tapping task produced robust activations in both the primary motor cortex (M1) and supplementary motor area (SMA) for each one of the five subjects. These results demonstrate that our novel concurrent TMS-fMRI setup allows us to detect activity in the dorsal surface of the brain. Note that the finger tapping periods were accompanied by visual cues (as prompts for the finger taps) and thus activations in both visual cortex and task-positive regions in the frontal and parietal regions can be detected for many subjects. For display purposes, the activation maps are generated using *p* < .001, uncorrected. Colors indicate t-values.

Further, we checked whether the position of the TMS coil remained stable over the course of the experiment. Using the anatomical images collected in the beginning and end of the experiment, we calculated the exact entry points of TMS stimulation based on vitamin E capsules placed on the TMS coil (**Figure 1C** and Methods). We then calculated the distance between the entry points in the beginning and end of the experiment and found that the average displacement across subjects was 2.32 mm (range: 1.9 - 2.8 mm). Therefore, there was only a modest shift in coil position (less than the length of a single voxel) from the beginning to the end of the experiment.

Finally, we investigated whether TMS led to subject movement that can contaminate our results. We found that the average framewise displacement (FD) (Power et al., 2012) was not different during periods of stimulation and periods of rest (t(4) = −2.24, *p* = 0.09). Similarly, the average FD was similar across the three TMS conditions (Low intensity 1 Hz: 1.05 mm, High intensity 1 Hz: 1.02 mm, High intensity bursts: 1.03 mm; F(1,2) = 0.05, *p* = 0.95). These results suggest that our main results below are unlikely to be caused by subject movement associated with TMS delivery.

#### No difference between high- and low-intensity TMS on local brain activity

Having validated our setup, we turned our attention to the question of whether TMS to DLPFC affects the local BOLD activity. To explore specifically the direct neural effects of TMS, we generated a contrast for High intensity 1 Hz > Low intensity 1 Hz, as well as a contrast for High intensity bursts > Low intensity 1 Hz. We found that both comparisons failed to show a clear increase around the targeted area even at the liberal threshold of *p* < .001 uncorrected (**Figure 3**). Specifically, subjects 1 and 2 showed hints of positive and negative local activations, respectively, whereas the rest of the subjects showed no positive or negative activations in the vicinity of the targeted area. Nevertheless, it could be argued that *p* < .001, uncorrected is still a fairly conservative threshold, which may mask the existence of small but consistent activations. Therefore, we examined the same brain activation maps at *p* < .05, uncorrected (**Supplementary Figure 2**) and found that the pattern of results remained the same.

**Figure 3.**
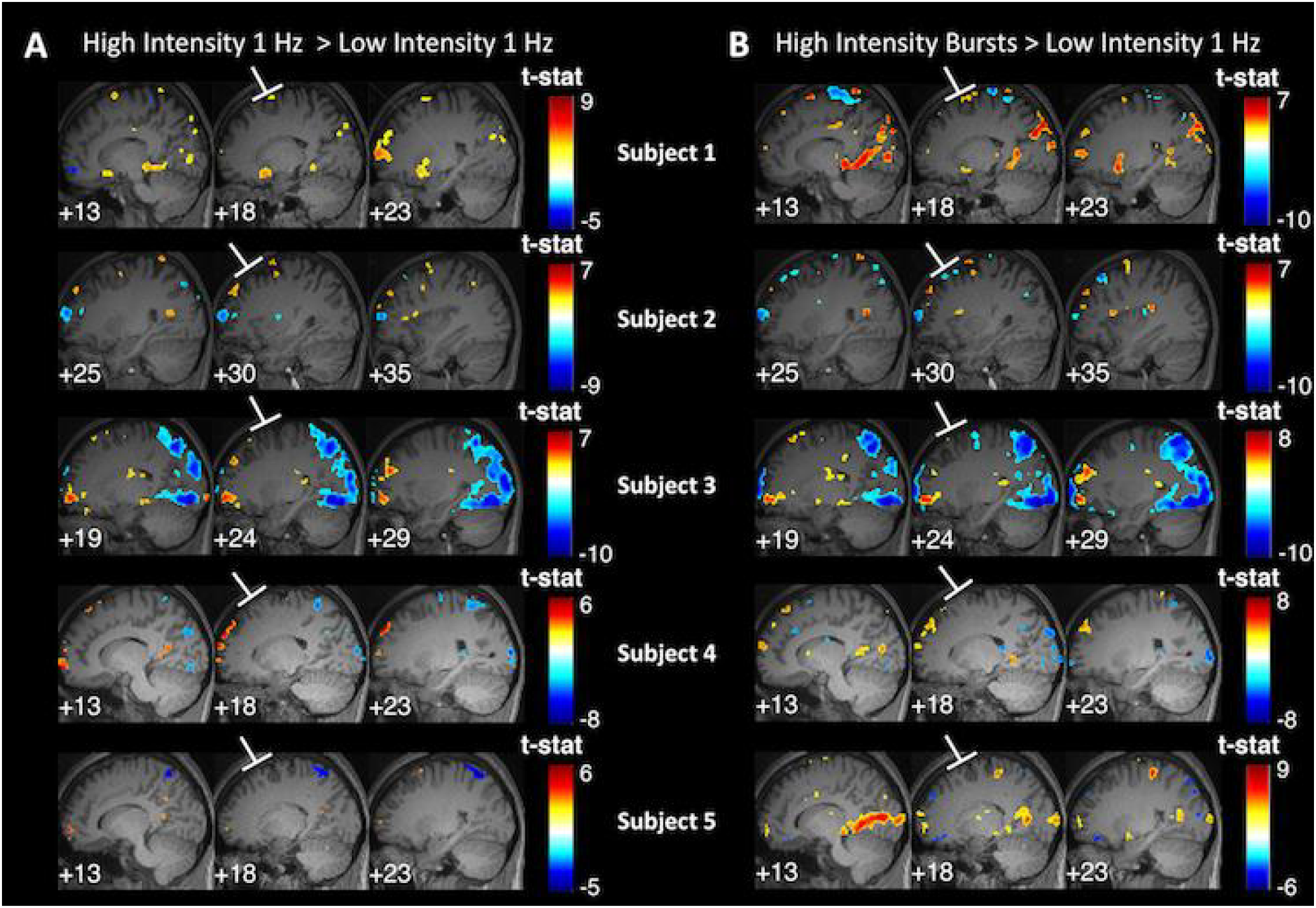
TMS to DLPFC produces no activation in the targeted area. We contrasted each of the two high-intensity TMS conditions against the low-intensity TMS condition: (A) High intensity 1 Hz > Low intensity 1 Hz contrast, and (B) High intensity bursts > Low intensity 1 Hz contrast. For both panels, the middle column shows slices centered at the site of stimulation, whereas the left and right columns show slices offset by 5 mm to the left and right from the slice in the middle column. The activation maps show a lack of systematic activation immediately in the targeted area and suggest the presence of large variability in remote areas across subjects. Activation maps generated using *p* < .001, uncorrected; colors indicate t-values.

To further explore the effect of TMS specifically on the targeted area, we defined spherical ROIs with radii of 8, 12, 16, and 20 mm at the location of stimulation and examined the BOLD signal change in these ROIs. We found no consistent pattern of positive or negative activations for any of the contrasts between the three different TMS conditions (**Figure 4**). Specifically, the two contrasts comparing high- to low-intensity TMS (“High intensity 1 Hz > Low intensity 1 Hz” and “High intensity bursts > Low intensity 1 Hz”) featured a total of 20 positive and 20 negative contrast estimates. Overall, the parameter estimates were not significantly different from zero for any of the 12 comparisons (3 contrasts × 4 ROI sizes; all *p*’s > .2). Finally, when examining each individual subject separately, we found that a total of three out of the possible 60 comparisons (3 contrasts × 4 ROI sizes × 5 subjects) were significantly higher than zero at the .05 level. This number of significantly positive activations (3/60 = 5% of all comparisons) is similar to what would be expected to happen by chance in the absence of any true differences between the conditions. These results show that there was no reliable difference in the level of local BOLD signal change between the three TMS conditions despite the large differences in stimulation intensity and number of pulses.

**Figure 4.**
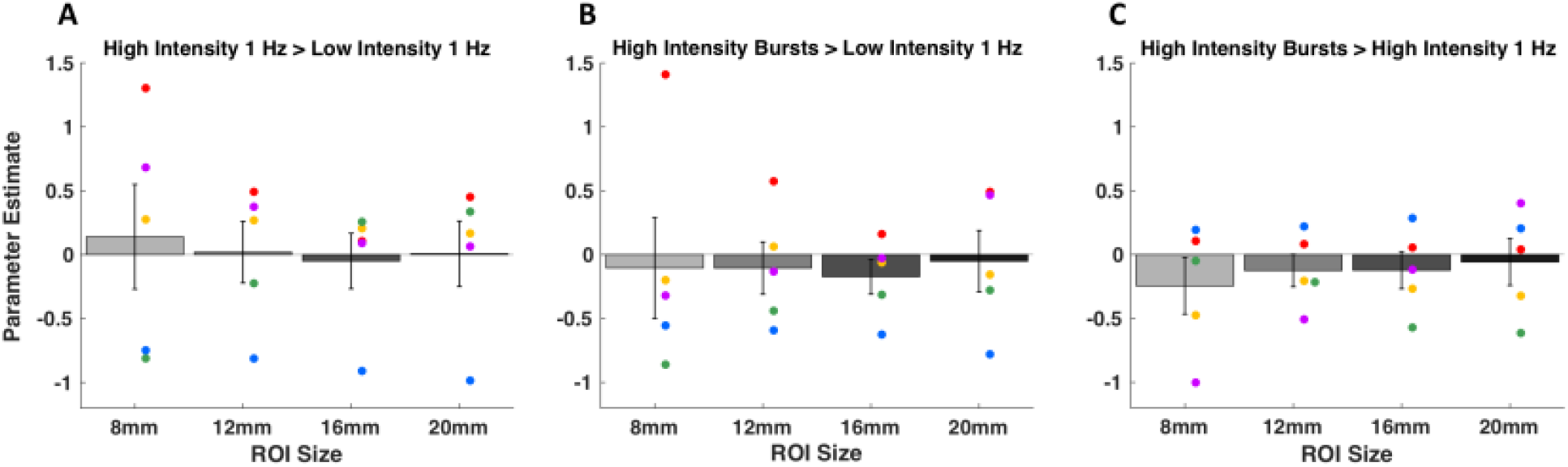
Parameter estimates for the comparisons between the three different TMS conditions in ROIs of different sizes. We plotted the parameter estimates for each contrast in the direction of the expected activations (high minus low intensity and more pulses minus fewer pulses). We examined ROIs defined at the location of stimulation of four different sizes using radii of 8, 12, 16, and 20 mm. (A) High intensity 1 Hz > Low intensity 1 Hz contrast. (B) High intensity bursts > Low intensity 1 Hz contrast. (C) High intensity bursts > High intensity 1 Hz contrast. We found no consistent pattern of positive or negative activations for any of the three contrasts and any of the ROI sizes. Error bars represent SEM, colors represent unique subjects.

Finally, it is possible that these null results are driven by TMS having an effect on BOLD activity that is not well described by the standard HRF. To examine this possibility, we plotted the average time courses of activity for each TMS condition using a finite impulse response (FIR) analysis. The FIR analysis does not assume a specific HRF shape and hence could reveal differences in the time courses of BOLD responses in a model-free manner. The time courses were computed for the 8-mm ROI that most closely reflects the local brain activity at the area of stimulation but similar results were obtained for the other ROI sizes too. We found no significant difference in the three time courses (F(20,240) = 0.73, *p*_*GG*_ = 0.64; **Figure 5**), suggesting that the lack of difference between the three TMS conditions in the GLM analyses is not driven by the assumed shape of the HRF.

**Figure 5.**
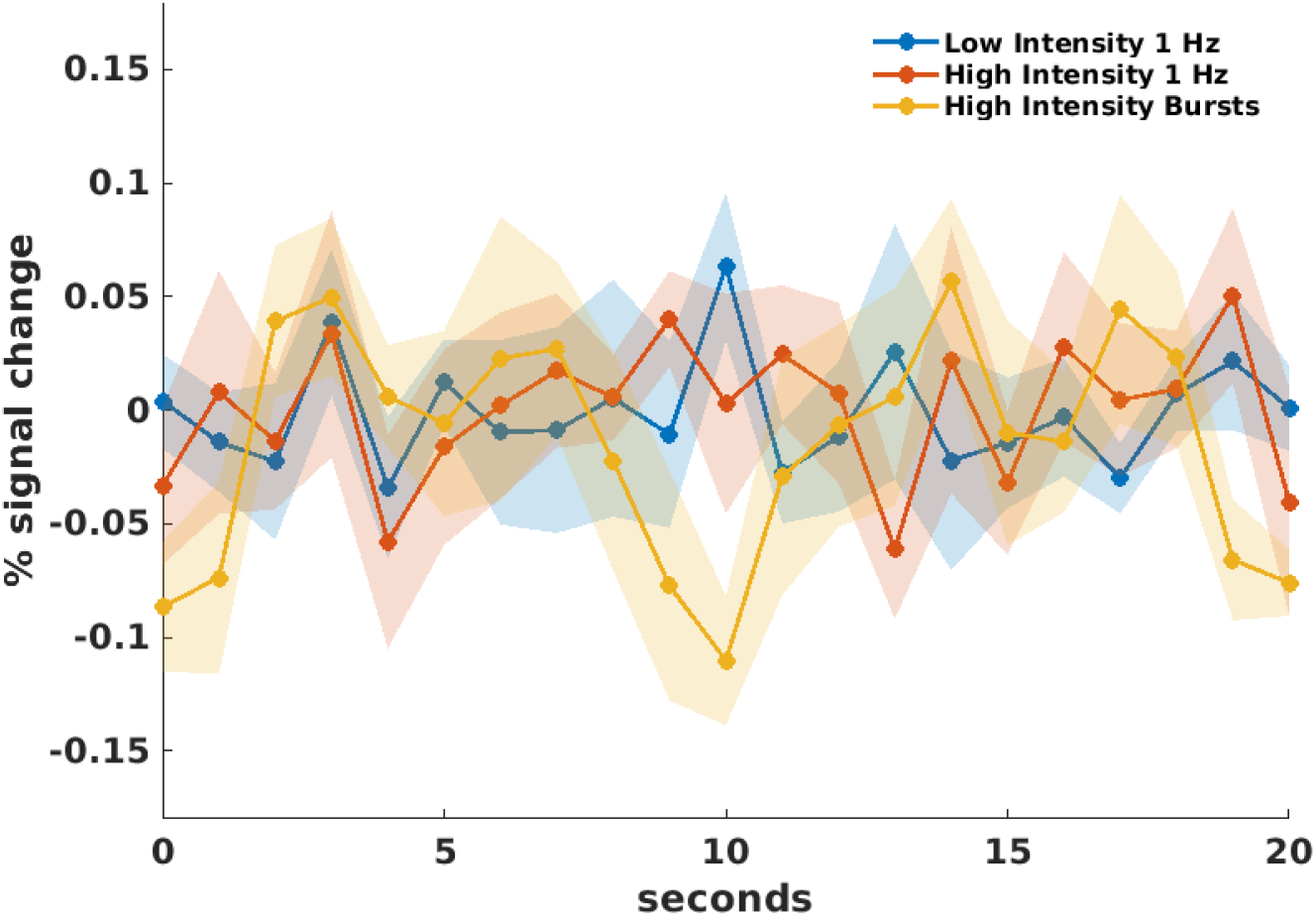
Finite impulse response (FIR) analysis. The figure plots the percentage of signal change as a function of time based on an FIR analysis in the 8-mm spherical ROI defined at the area of stimulation. There was no significant difference between the overall time courses for the three conditions. Each data point shows the average percentage of signal change for all subjects and the shadowed areas depict s.e.m.

#### TMS condition can be decoded at the area of stimulation

The lack of consistent increase in local BOLD activity raises the question of whether any TMS effects at the targeted area can be detected using fMRI at all. To address this question, we examined the distributed patterns of activity and built an MVPA classifier which attempted to distinguish the different TMS conditions. We built a separate classifier for each of the four ROI sizes (radii of 8, 12, 16, and 20 mm) and used 3-fold cross validation by training the classifier on two TMS runs and testing on the remaining run.

We first performed 3-way classification on the three TMS conditions (Low intensity 1 Hz, High intensity 1 Hz, and High intensity bursts). Across the four ROI sizes, 17 out of the 20 classifications were above chance level, two were below chance level, and one was exactly at chance level. To quantify the classification accuracy for each ROI, we performed a mixed model analysis on the individual TMS trials for classification with subject as a random effect and accuracy of classification as the dependent variable. We found that the intercept value indicating the classification accuracy in the group was significantly higher than the intercept expected by chance-level classification for three of the four ROI sizes (8-mm ROI: t(449) = 4.58, *p* = 6 × 10^−6^; 12-mm ROI: t(449) = 1.95, *p* = 0.0519; 16-mm ROI: t(449) = 2.36, *p* = 0.0186; 20-mm ROI: t(449) = 7.24, *p* = 2 × 10^−12^; **Figure 6A**). The larger ROI sizes generally produced higher intercepts indicating better classification performance, likely due to the larger number of voxels included. These results indicate that MVPA could successfully distinguish the BOLD activity patterns evoked by the different TMS conditions despite the absence of consistent univariate activations.

**Figure 6.**
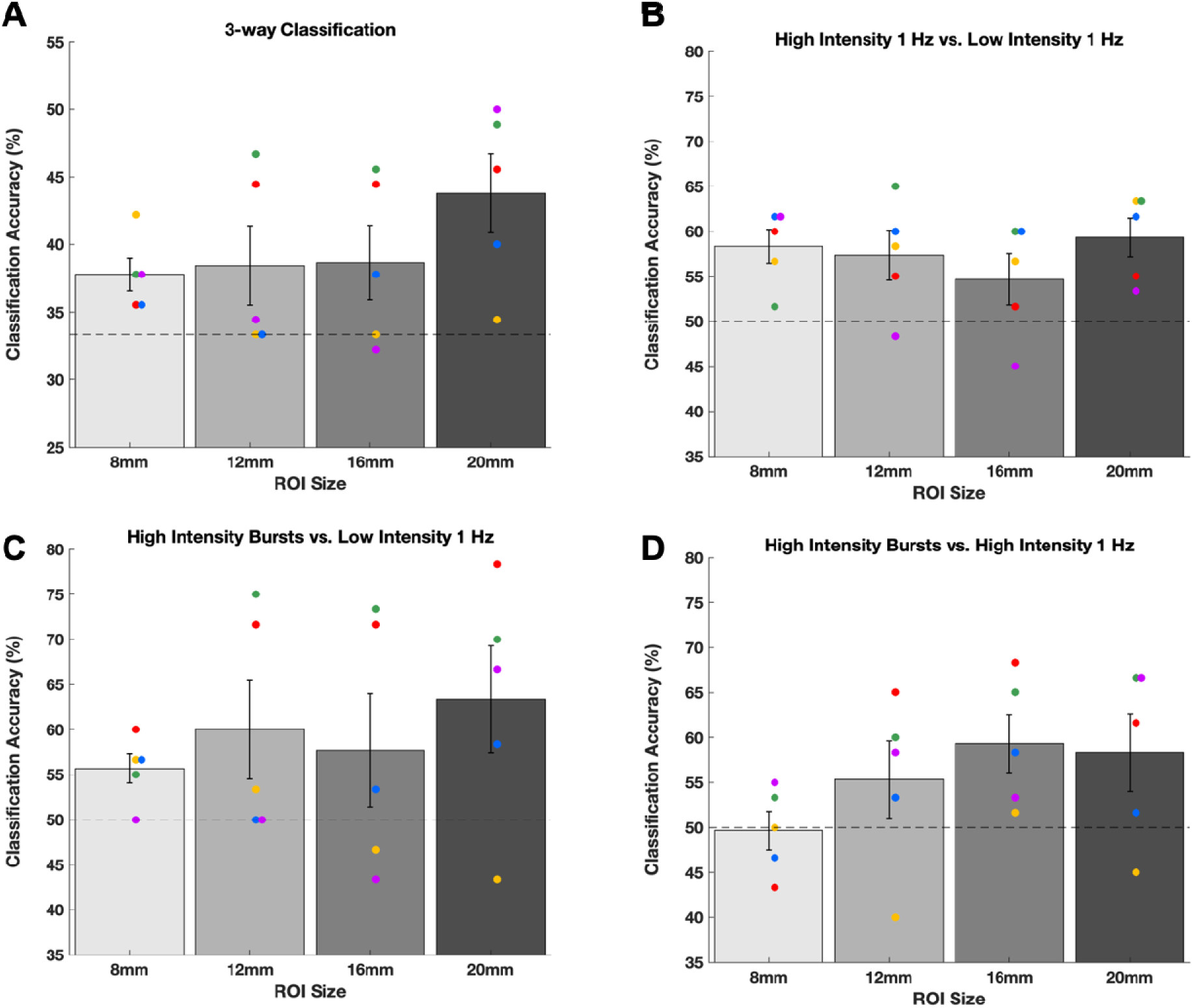
Decoding the TMS condition using MVPA in different ROI sizes at the area of stimulation. (A) Three-way classification performance for the different TMS conditions (High intensity bursts, High intensity 1 Hz, and Low intensity 1 Hz). (B-D) Two-way classification performance for each pairwise comparison of different TMS conditions. Dashed lines show chance level (33.33% in Panel A, 50% in Panels B-D). Error bars represent SEM, colors represent unique subjects.

Further, we examined whether the above-chance classification performance in the 3-way classification was due to any specific pair of conditions. To do so, we build additional 2-way MVPA classifiers to distinguish between all pairs of conditions drawn from the three TMS conditions. We first investigated classification performance in the 20-mm ROI since that ROI provided the best 3-way classification performance. We found that the classification accuracy of the combined data using mixed model analysis was numerically highest when comparing the High intensity bursts vs. Low intensity 1 Hz conditions (accuracy = 63.3%, chance level = 50%, t(299) = 3.37, *p* = 8 × 10^−4^; **Figure 6C**) and slightly lower for the other two comparisons (High intensity 1 Hz vs. Low intensity 1 Hz: accuracy = 59.3%, t(299) = 5.58, *p* = 6 × 10^−8^; High intensity bursts vs. High intensity 1 Hz: accuracy = 58.3%, t(299) = 5.56, *p* = 6 × 10^−8^; **Figure 6B,D**) but the difference in classification accuracy was not significantly different between the three comparisons (all three *p*’s > .1, Z tests for proportions). We found qualitatively similar results for the 16-mm, 12-mm, and 8-mm ROIs (**Figure 6B-D**) with classification accuracy generally being above chance but not significantly different between the three comparisons. These results suggest that the successful 3-way classification was likely not driven by any one particular condition.

### Experiment 2

Experiment 1 demonstrated that variations in the intensity of stimulation and number of pulses delivered do not result in local activity changes but can be successfully distinguished using MVPA. In Experiment 2, we sought to extend these findings by exploring whether the same pattern of results would be obtained when varying TMS frequency while keeping both the intensity of stimulation (100% of motor threshold) and number of pulses (30 per train) the same across conditions. We again targeted DLPFC but used four different frequencies – 5, 8.33, 12.5, and 25 Hz – and analyzed the pattern of activity at the area of stimulation by employing both univariate and multivariate techniques.

Similar to Experiment 1, we first computed TMS coil displacement from the beginning to the end of the experiment, and found that the movement of the coil was relatively small (average = 2.23 mm, range: 1-3.7 mm). Further, the average subject movement was not significantly different during stimulation and rest periods (t(11) = 0.04, *p* = 0.97) or during different TMS conditions (5 Hz stimulation: 1.246 mm, 8.33 Hz stimulation: 1.597, 12.5 Hz stimulation: 1.853 mm, 25 Hz stimulation: 1.385 mm; F(1,3) = 0.83, *p* = 0.48), suggesting that TMS did not result in increased subject movement.

#### Stimulation frequency does not modulate local BOLD activity

We first compared the BOLD signal change between every two pairs of frequencies. We found no reliable univariate effects: no frequency produced consistently higher activity in the vicinity of the targeted area compared to any other frequency even at the liberal threshold of *p* < .001 uncorrected (**Figure 7**). Similar to Experiment 1, we observed substantial across-subject variability in the activations induced in remote areas but relatively smaller within-subject variability across the two sessions completed by each subject.

**Figure 7.**
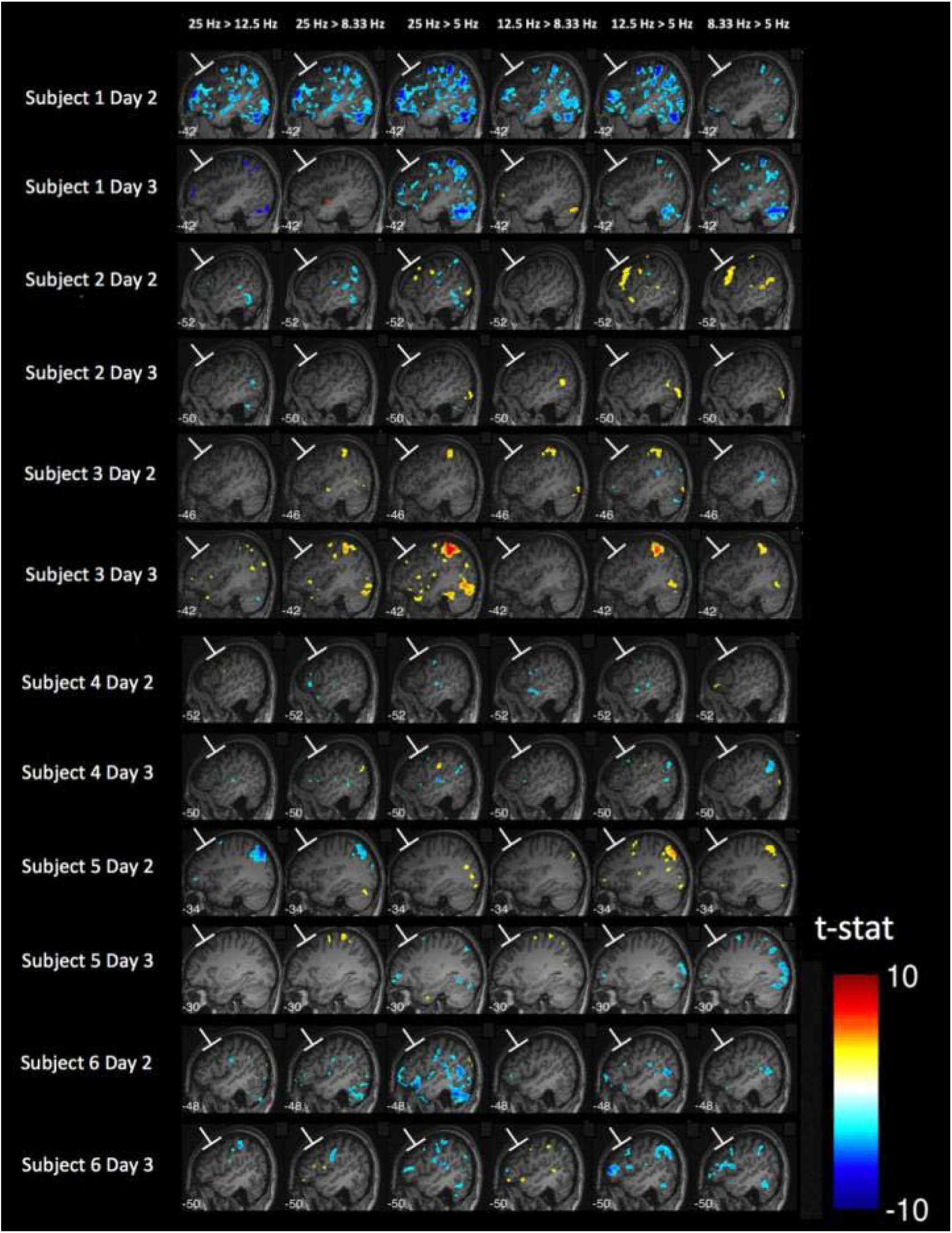
Different TMS frequencies do not produce differential activation in the vicinity of the targeted area. Sagittal slices of the brain are shown at the location of stimulation spot for each of the six pairwise comparisons between the four different TMS frequencies used in Experiment 2. For each subject, data from the two TMS sessions (Day 2 and Day 3) are shown in separate rows. No frequency resulted in consistently higher activity than any other frequency. Activation maps generated using *p* < .001, uncorrected; colors indicate t-values.

As in Experiment 1, we further explored the local activity by examining the BOLD effects in spherical ROIs with radii of 8, 12, 16, and 20 mm. We again found no consistent pattern of positive or negative activations for any of the contrasts between the four different TMS frequencies (**Figure 8**). We performed mixed model analyses on the individual-subject parameter estimates with subject and day of testing as random effects. We found that the intercept for the parameter estimates were not significantly different from zero for any of the 24 comparisons (6 pairs of conditions × 4 ROI sizes; all *p*’s > .05). Finally, when examining each comparison for each individual subject separately, we found that a total of 12 out of the possible 288 comparisons (6 pairs of conditions × 4 ROI sizes × 6 subjects × 2 days) were significantly different than zero at the .05 level (rate of significance = 4.17%, which is similar to the 5% rate that would be expected by chance). These results suggest that the four different TMS frequencies did not result in reliable differences in local BOLD activity.

**Figure 8.**
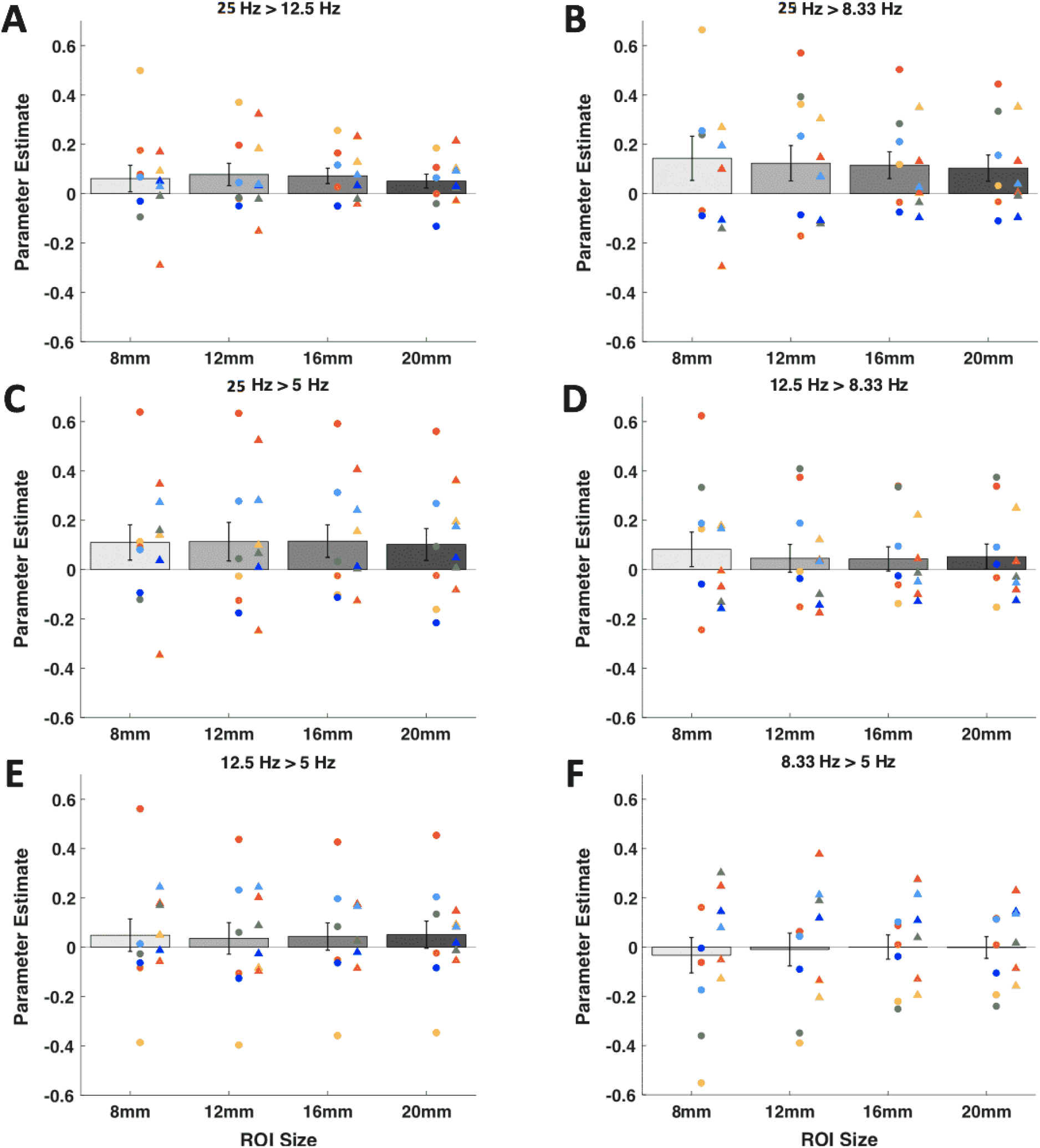
Parameter estimates for the comparisons between the four different TMS frequencies in ROIs of different sizes. We plotted the parameter estimates for higher minus lower TMS frequencies. We further examined ROIs defined at the area of stimulation of four different sizes using radii of 8, 12, 16, and 20 mm. (A) 20 Hz > 12.5 Hz contrast. (B) 20 Hz > 8.33 Hz contrast. (C) 20 Hz > 5 Hz contrast. (D) 12.5 Hz > 8.33 Hz contrast. (E) 12.5 Hz > 5 Hz contrast. (F) 8.33 Hz > 5 Hz contrast. We found no consistent pattern of positive or negative activations for any of the six contrasts and any of the ROI sizes. Error bars represent SEM, colors represent unique subjects with data from Day 2 plotted as a circle and data from Day 3 plotted as a diamond.

Beyond comparing the four TMS conditions, we also examined whether we could observe any activations related to the presence of TMS pulses. We conducted a group-level analysis and, as expected, found consistent activations in bilateral auditory cortex that is likely caused by the clicking sounds produced by the TMS pulses (**Supplementary Figure 3**). We note that we could not perform a similar analysis for Experiment 1 because we did not have full brain coverage there. Finally, as in Experiment 1, we also performed an FIR analysis and found no significant differences between the time courses for the four conditions (**Supplementary Figure 4**), indicating that the lack of significant differences between the conditions is not due to the assumed HRF shape.

#### The frequency of stimulation can be decoded at the area of stimulation

Similar to Experiment 1, we examined whether the lack of consistent local BOLD activity differences between the four TMS frequencies implies that TMS frequency has no impact on local BOLD activity at all. To address this question, we again built an MVPA classifier to distinguish between the different TMS frequencies. A separate classifier was constructed for each of the four ROI sizes (radii of 8, 12, 16, and 20 mm). We again used 3-fold cross validation by training on two TMS runs and testing on the remaining run, and performed a 4-way classification on the four frequencies.

Performance of all 48 classifiers (6 subjects × 2 days × 4 ROI sizes) was above chance level. As in Experiment 1, to quantify the classification accuracy for each ROI, we performed mixed model analyses with subject and day of testing as random effects and individual trial accuracy as the dependent variable. We found that the intercept was significantly higher than expected by chance for all ROI sizes (8-mm ROI: t(708) = 20.77, *p* = 3.4 × 10^−75^; 12-mm ROI: t(708) = 24.33, *p* = 1.7 × 10^−95^; 16-mm ROI: t(708) = 24.54, *p* = 9.8 × 10^−97^; 20-mm ROI: t(708) = 22.60, *p* = 1.4 × 10^−86^; **Figure 9**). The classification accuracy increased slightly for larger ROI sizes possibly because they contain more voxels compared to smaller ROIs. These results show that, just as in Experiment 1, decoding analyses based on the pattern of BOLD activations at the area of stimulation successfully distinguished the four different TMS frequencies even in the absence of consistent univariate differences between these frequencies.

**Figure 9.**
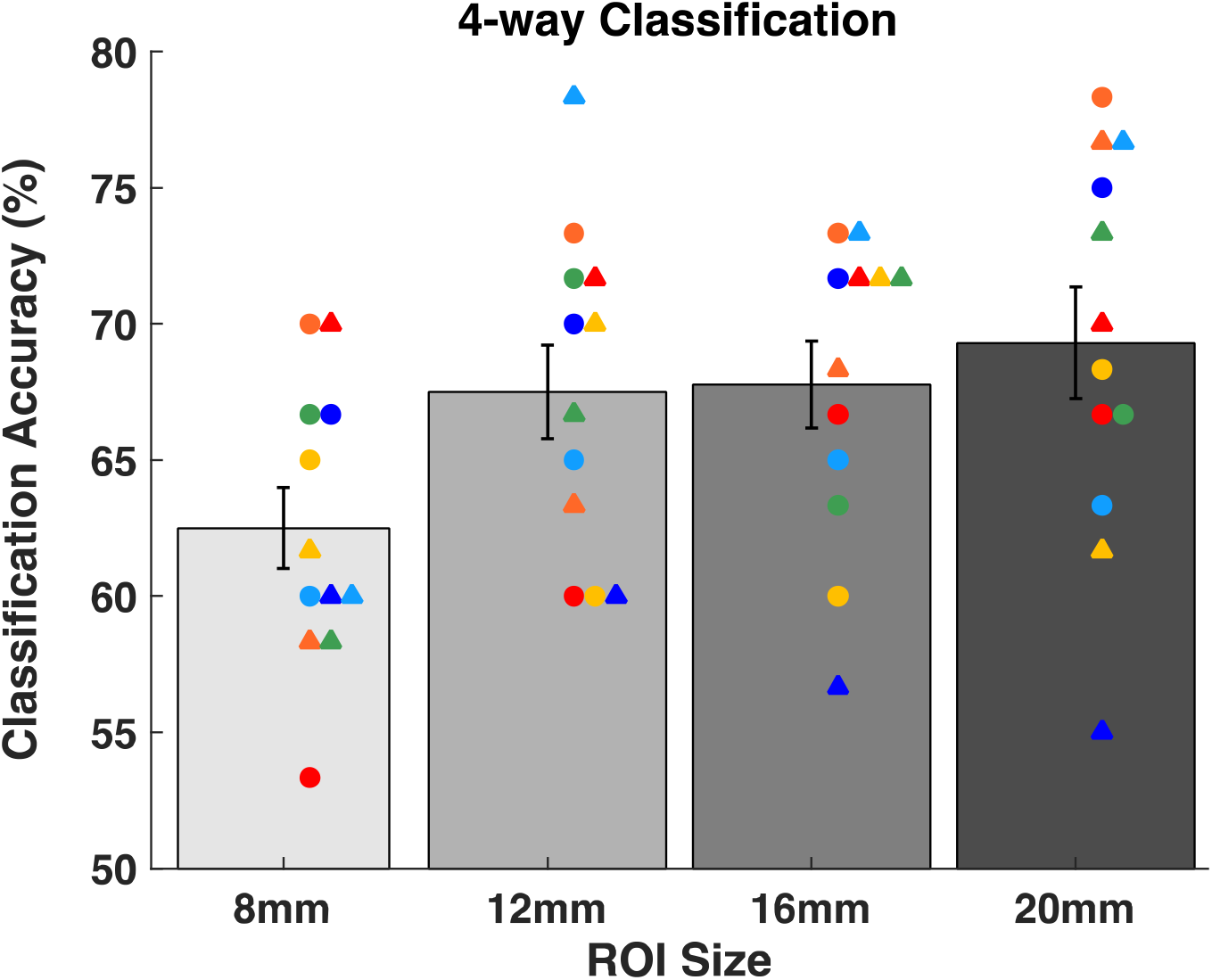
Decoding the TMS frequency using MVPA in different ROI sizes at the area of stimulation. Four-way classification performance for the four different TMS frequencies (5, 8.33, 12.5, and 25 Hz). Significant classification accuracy was obtained for all ROI sizes. Error bars represent SEM, colors represent unique subjects with data from Day 2 plotted as circles and data from Day 3 plotted as triangles. Chance level classification is 25%.

We further examined the relationship between the different frequencies by testing whether a classifier trained on one pair of frequencies can be used to classify a different pair of frequencies. Specifically, we trained a classifier to distinguish between 5 and 25 Hz TMS using the voxels in the 20-mm ROI. This classifier had very high training accuracy (mean classification accuracy = 97.50%, t(348) = 57.65, *p* = 4.1 × 10^−180^; **Figure 10**) demonstrating that the 5 and 25 Hz conditions resulted in very different patterns of activity despite the fact that neither led to consistently higher activations than the other. Critically, we used this same classifier to discriminate between 8.33 and 12.5 Hz TMS. We found that the classifier trained on 5 vs. 25 Hz stimulation was able to reliably distinguish between 8.33 and 12.5 Hz stimulation (mean classification accuracy = 80.28%, t(348) = 14.42, *p* = 2.8 × 10^−37^; **Figure 10**), suggesting that the activation patterns changed monotonically for the four frequencies.

**Figure 10.**
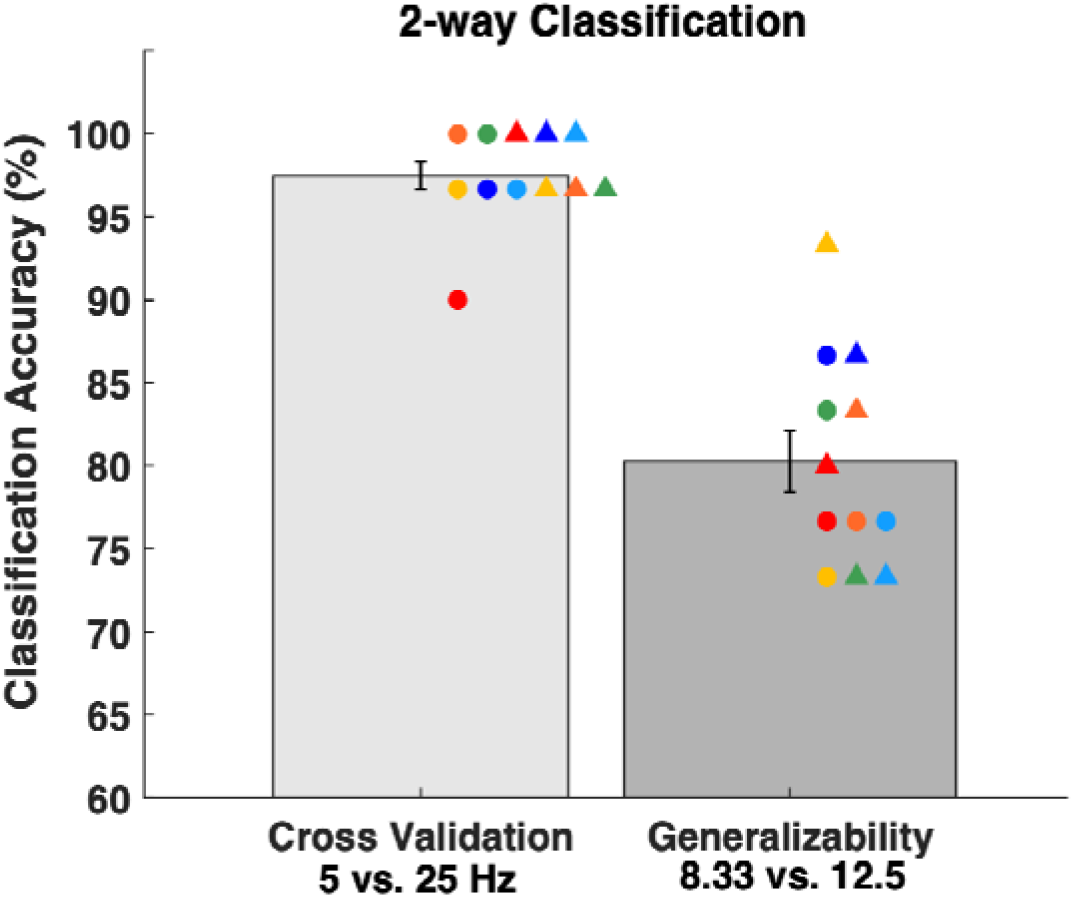
Relationship between the activation patterns of different frequencies. We trained a classifier to distinguish between 5 and 25 Hz stimulation and tested whether it can distinguish between 8.33 and 12.5 Hz stimulation. The classifier had very high cross-validation accuracy for the 5 vs. 25 Hz classification. Critically, it was able to also distinguish between 8.33 and 12.5 Hz stimulation even though it was never trained on these frequencies. Chance level is 50%.

One possible reason for the high decoding performance at the targeted area in absence of univariate activation is that the decoded information content could stem from experimental factors other than direct neuronal effects of TMS. For example, TMS delivery may have non-specific effects such as induced subject movement (though, as we showed above, average movement was matched across conditions). To ensure that the above-chance decoding performance was not due to such effects, we performed a control analysis in which we defined 20-mm ROIs in left and right visual cortices and assessed the classification performance in these ROIs. We found that the classification performance at the stimulation spot was significantly greater than the classification performance in both left visual cortex (t(708) = 10.06, *p* = 2.3 × 10^−22^) and right visual cortex (t(708) = 11.52, *p* = 2.99 × 10^−28^; **Supplementary Figure 5**). These results show that the high classification performance at the spot of stimulation is unlikely to be purely due to non-specific effects of TMS since these effects would be comparable in the visual cortex ROIs. On the other hand, MVPA analyses in ROIs defined in the left and right auditory cortex showed high classification performance (**Supplementary Figure 5**), presumably due to the different sound characteristics associated with the TMS delivery in each condition.

Finally, we examined whether the voxels that were most informative for classification were located near the surface of the brain. Given the rapid decrease of the magnetic field away from the TMS coil, we hypothesized that TMS should have a smaller influence on the voxels in the lower half of the large ROI sizes. To test this hypothesis, we performed dimensionality reduction where we identified the most informative voxels (that is, the voxels with the most information with respect to classes; Langs et al., 2011). We found that the majority (63.92%) of these voxels were located in the upper half of the 20-mm ROI where the TMS magnetic pulses are the strongest (t(11) = 13.92, p = 0.0007). This clustering of informative voxels in the upper half of 20-mm ROI occurred despite the fact that the number of gray matter voxels in the upper vs. lower half of the ROI were not significantly different (t(11) = 0.68, p = 0.5134). These results demonstrate that the voxels closest to the surface of the brain carried the most information about classification, consistent with the known decrease of magnetic strength with distance.

## Discussion

We investigated the effect of TMS on BOLD activity and MVPA decodability at the targeted area. In two experiments, we observed that both the intensity and frequency of stimulation could be reliably decoded from local brain activity. On the other hand, TMS had no effect on overall BOLD activity at the area of stimulation even when bursts of up to 30 high-intensity pulses were employed. These results suggest that the effects of TMS on the targeted brain area may be more complex than previously appreciated but that decoding methods are sensitive to variations in both TMS intensity and frequency.

Many previous concurrent TMS-fMRI studies that examined local BOLD activity delivered TMS over motor (Bohning et al., 1999, 2000a; Baudewig et al., 2001; Bestmann et al., 2003, 2004; Denslow et al., 2005a) or visual cortex (Caparelli et al., 2010). However, a major difficulty in interpreting the results of these studies is that the observed activations could be a combination of local effects and afferent feedback arising from the induced twitches or visual phosphenes (Bestmann et al., 2008). Therefore, in order to specifically examine the local effects of TMS independent of overt behavioral responses such as twitches and phosphenes, here we delivered TMS over DLPFC and examined its effect on the voxels located immediately at the area of stimulation.

Why did TMS not increase local BOLD activity in our study? This can be seen as puzzling given that TMS is known to induce action potential at the area of stimulation, which should then drive increased BOLD activity. There are a number of possible reasons. First, a number of studies demonstrated that TMS pulses can lead to an immediate increase followed by a prolonged decrease in the firing rate of individual neurons in rats (Li et al., 2017), cats (Moliadze et al., 2003; Kozyrev et al., 2014), and monkeys (Grigsby, 2015; Romero et al., 2019b). These findings imply that, when integrated over an extended period of time, overall neural activity following a TMS pulse may not actually show an overall increase. Given the slow nature of the BOLD signal (Jezzard et al., 2001; Faro and Mohamed, 2006), it is likely that the observed activity in fMRI studies reflects the overall neural firing over an extended period of time rather than just the initial burst of firing, and therefore these animal studies could explain the lack of change in the BOLD signal observed here. Second, TMS may preferentially affect the white matter tracts that leave an area, rather than the cell bodies in that area. If so, one may expect that the TMS effects should be seen in downstream areas of the brain but not locally at the area of stimulation. Finally, TMS may have additional effects other than making some neurons fire, and those effects may contribute to the BOLD response. For example, recent studies have suggested that TMS can directly inhibit cortical dendrites (Murphy et al., 2016). Note that these possibilities are not mutually exclusive and several of these processes may contribute to the fact that TMS does not produce reliable local activations. Testing these possibilities requires animal studies where neuronal spiking and BOLD activity are measured simultaneously. Beyond revealing more about the mechanisms of TMS, such studies can shed further light on the nature and correlates of the BOLD signal itself.

Our results add important new evidence regarding the local effects of TMS outside the motor and visual cortices in the absence of a task. Previous research on this topic found inconsistent results with some studies reporting an increase of activity in the vicinity of the targeted area (Nahas et al., 2001; Li et al., 2004a; Bestmann et al., 2005; Vink et al., 2018a), while many others reported no such increase (Baudewig et al., 2001; Kemna and Gembris, 2003; Li et al., 2004b; Sack et al., 2007; De Weijer et al., 2014; Hawco et al., 2017; Dowdle et al., 2018). Critically, the studies that reported increased activity in the targeted area in the absence of a task generally did not precisely estimate the exact location of the TMS coil for each subject. Given that TMS induces both sensory (auditory and tactile) stimulation and corresponding cognitive processes, it is possible that at least in some cases the reported activations were due to such processes and that the actual activation was not located exactly at the site of stimulation. Future studies should adopt an approach similar to the one taken here by precisely localizing the location of stimulation for each subject and conducting ROI analyses.

It is also possible that the differences between the studies that found local activations and studies that did not find such activations are due to differences in stimulation intensities, frequencies, number of pulses, or sites of stimulation. For example, it is possible that some sites of stimulation (e.g., premotor cortex) behave differently from others (e.g., DLPFC) or that local activations would only be observed for specific intensities or frequencies. Nevertheless, many of the studies that reported local activation used lower intensities and lower frequencies than other studies that found no local activations in roughly the same brain regions. Therefore, while the differences in stimulation parameters and locations could be a contributing factor, they are unlikely to account for all of the differences observed in the literature.

It should be noted, that our conclusions regarding a lack of local BOLD activity apply exclusively to the effects of TMS in the absence of a task. It is quite possible that when TMS is delivered during a task that recruits a specific brain region, TMS would have an overall univariate effect on the BOLD activity in that region. Indeed, some previous studies have used an ROI approach and found that TMS changed the local activity in the presence of a task (Feredoes et al., 2011; Heinen et al., 2014b) while simultaneously finding no effect in the absence of a task (Heinen et al., 2014b).

Our results showed that even though TMS did not affect the univariate local BOLD activity, the different TMS conditions could be reliably decoded at the targeted area using MVPA. Furthermore, we found that the most informative voxels for classification were located near the surface of the brain where the stimulation strength is the highest. These findings suggest that TMS does influence BOLD activity even in the absence of a task but that the TMS effects may differ even for nearby neuronal populations thus producing a complex pattern of activations and deactivations that can be detected using MVPA but not with standard univariate techniques. Such differences among neuronal populations could be due to variability in the predominant neuronal types, the properties of local network connectivity, the orientation of the cortical column relative to the magnetic field, etc. Future research in animals can test these possibilities more directly.

Finally, we note several limitations in our experiments. First, both of our experiments had only a few subjects. However, we collected large amount of data for each subject and performed the analyses at the individual level. This strategy was necessitated by the variability in the actual stimulation site between subjects, which does not allow for aggregation across subjects. Given that we observed similar results across our five subjects in Experiment 1 and all 12 individual sessions in Experiment 2, it is unlikely that larger samples would lead to different conclusions. In addition, we collected two different TMS sessions in Experiment 2 in order to increase the power to detect effects at the individual level. However, despite the use of neuronavigation, the actual site of stimulation differed slightly between the two sessions, making it impossible to pool the data across the two sessions as the exact area of stimulation had shifted. Therefore, to avoid such issues, we suggest that future concurrent TMS-fMRI experiments employ a single TMS session.

In conclusion, we found that, across two different experiments, TMS intensity and frequency could be decoded using MVPA even though TMS did not increase BOLD activity at the targeted area. These findings help settle a longstanding discrepancy in the literature and provide important insight into the mechanisms through which TMS influences neural activity.

## Acknowledgements

This work was funded by the National Institute of Health (award R21MH122825 to D.R.).

## Supplementary Figures

**Supplementary Figure 1.**
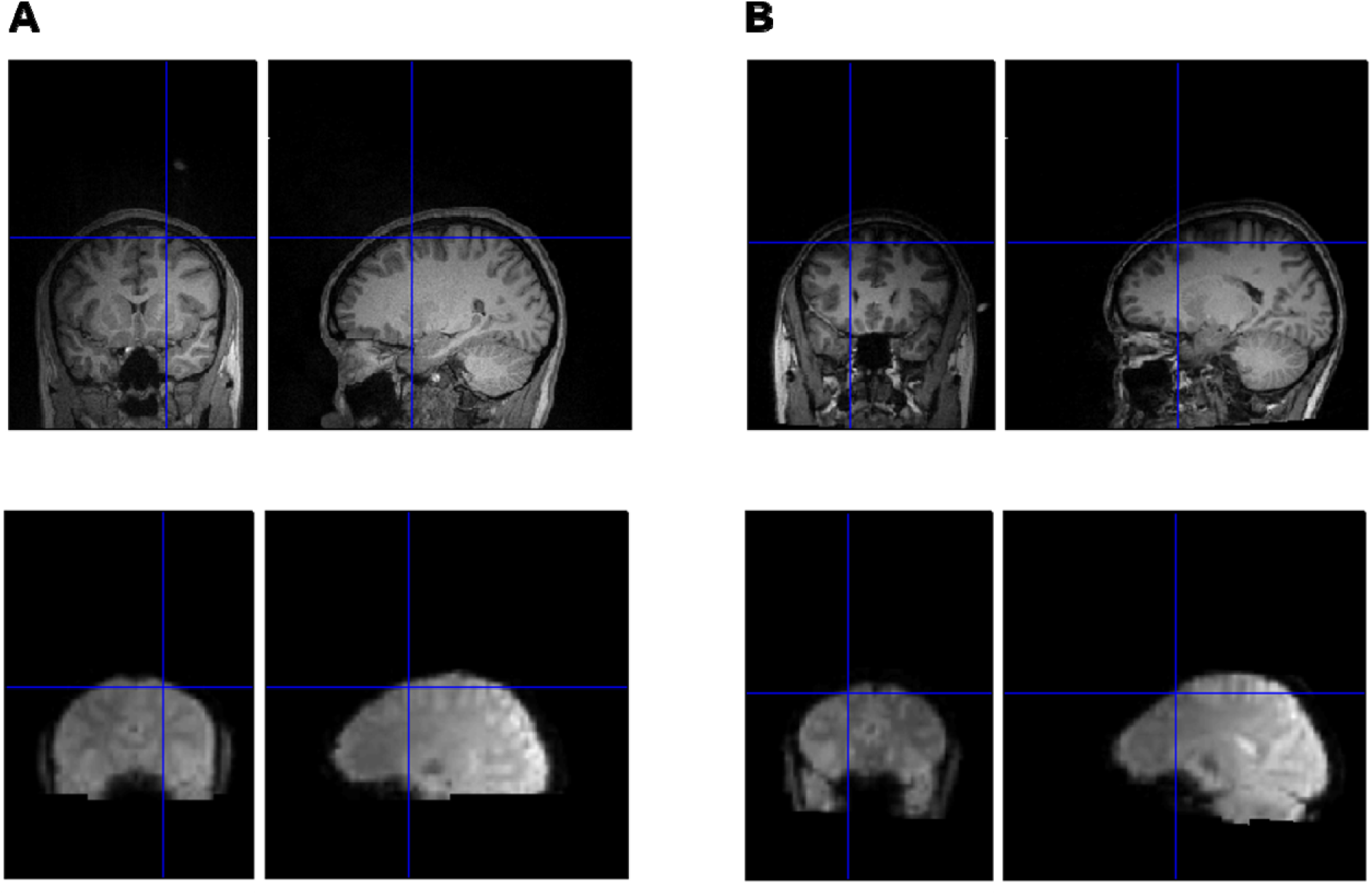
Quality of alignment of functional and anatomical images. The blue cross in all images shows the stimulation spot for a representative subject from (A) Experiment 1 and (B) Experiment 2. The top row shows the anatomical image of the subject, whereas the bottom row shows one volume from the functional scans. The quality of alignment was checked for all of the subjects visually using SPM and no systematic deviation was found. The figure also shows that there was no visible dropout or distortion in the functional images at the site of stimulation that could have been caused by the presence of the TMS coil.

**Supplementary Figure 2.**
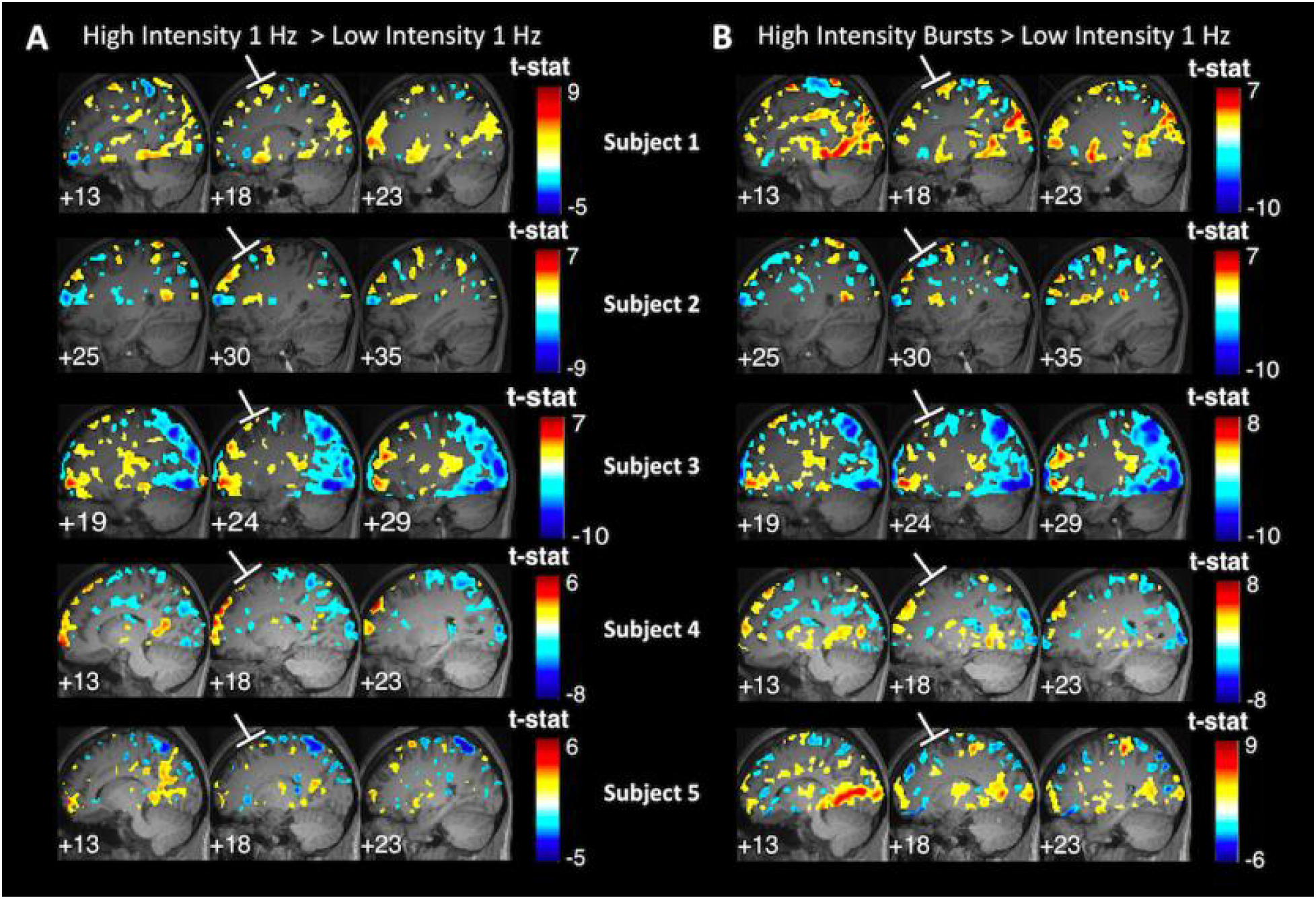
TMS to DLPFC produces no activation in the vicinity of the targeted area even at p < .05 uncorrected. We contrasted each of the two high-intensity TMS conditions against the low-intensity TMS condition, but, unlike in Figure 3, used an even more liberal threshold for significance (*p* < .05, uncorrected). (A) High intensity 1 Hz > Low intensity 1 Hz contrast. (B) High intensity bursts > Low intensity 1 Hz contrast. The activation maps show a lack of systematic activation in the vicinity of the targeted area and suggest the presence of large variability in remote areas across subjects. All figure details are as in Figure 3.

**Supplementary Figure 3.**
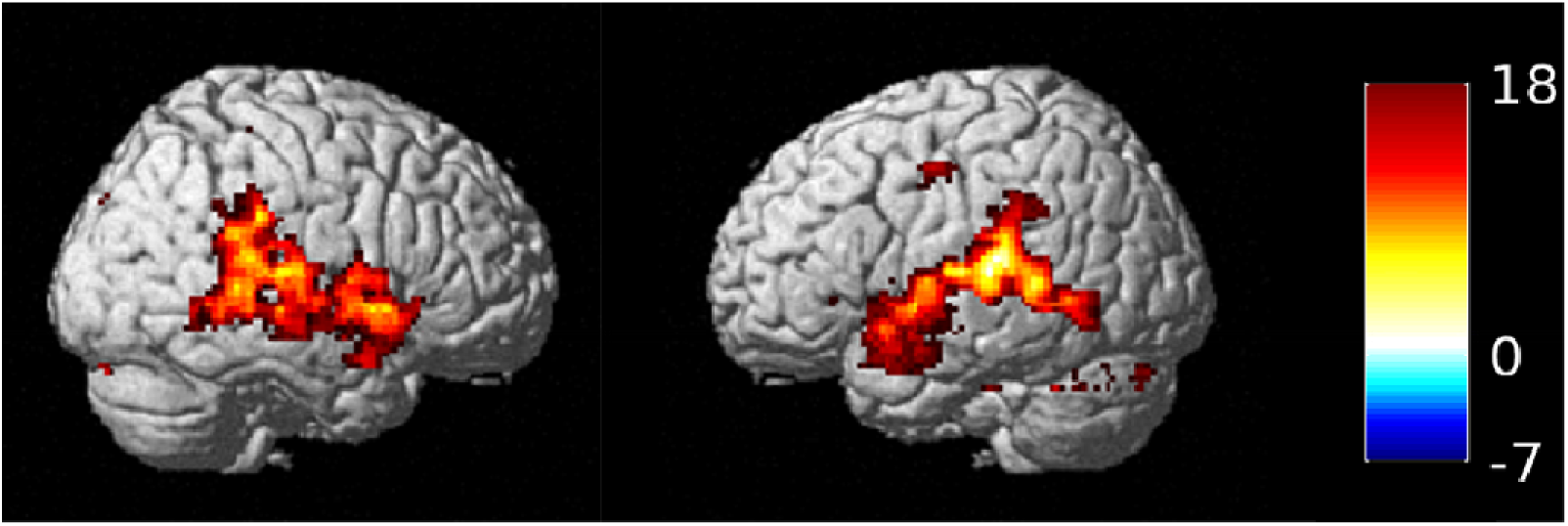
Univariate whole brain analysis in Experiment 2. A whole brain, second-level analysis on the normalized data from all subjects using the contrast TMS > baseline revealed consistent TMS-related activations in auditory cortices in Experiment 2. These activations were likely due to the click sound produced by TMS while stimulation. We note that similar analyses could not be run in Experiment 1 because we did not have whole brain coverage. For display purposes, the figure shows activations at *p* < 0.001, uncorrected.

**Supplementary Figure 4.**
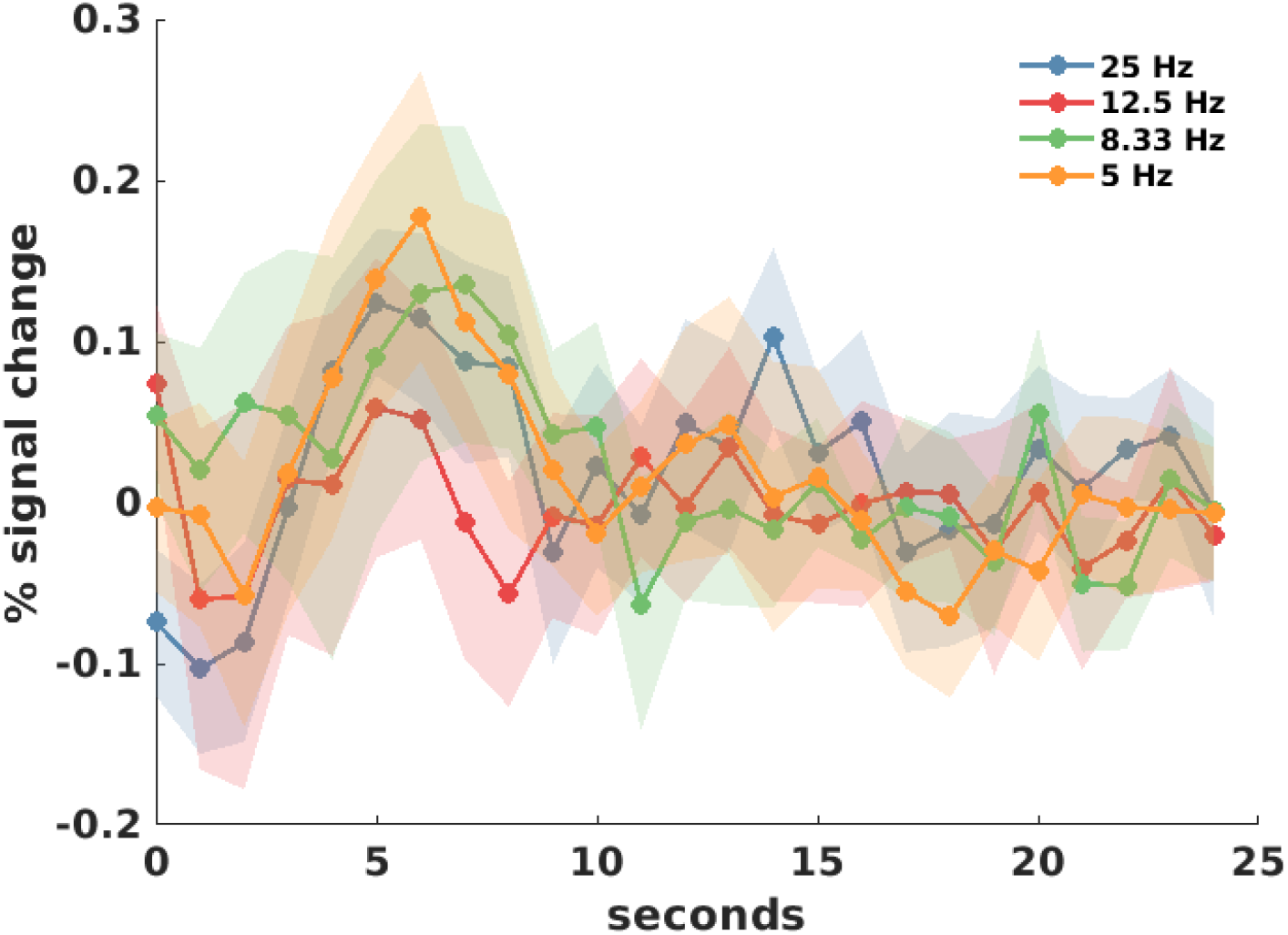
FIR analysis in Experiment 2. The figure plots the percentage of signal change as a function of time based on an FIR analysis in the 8-mm spherical ROI defined at the area of stimulation. There was no significant difference between the overall time courses for the four conditions at any time point. Each data point shows the average percentage of signal change for all subjects. Zero is the time of stimulus onset and the shadowed areas depict s.e.m.

**Supplementary Figure 5.**
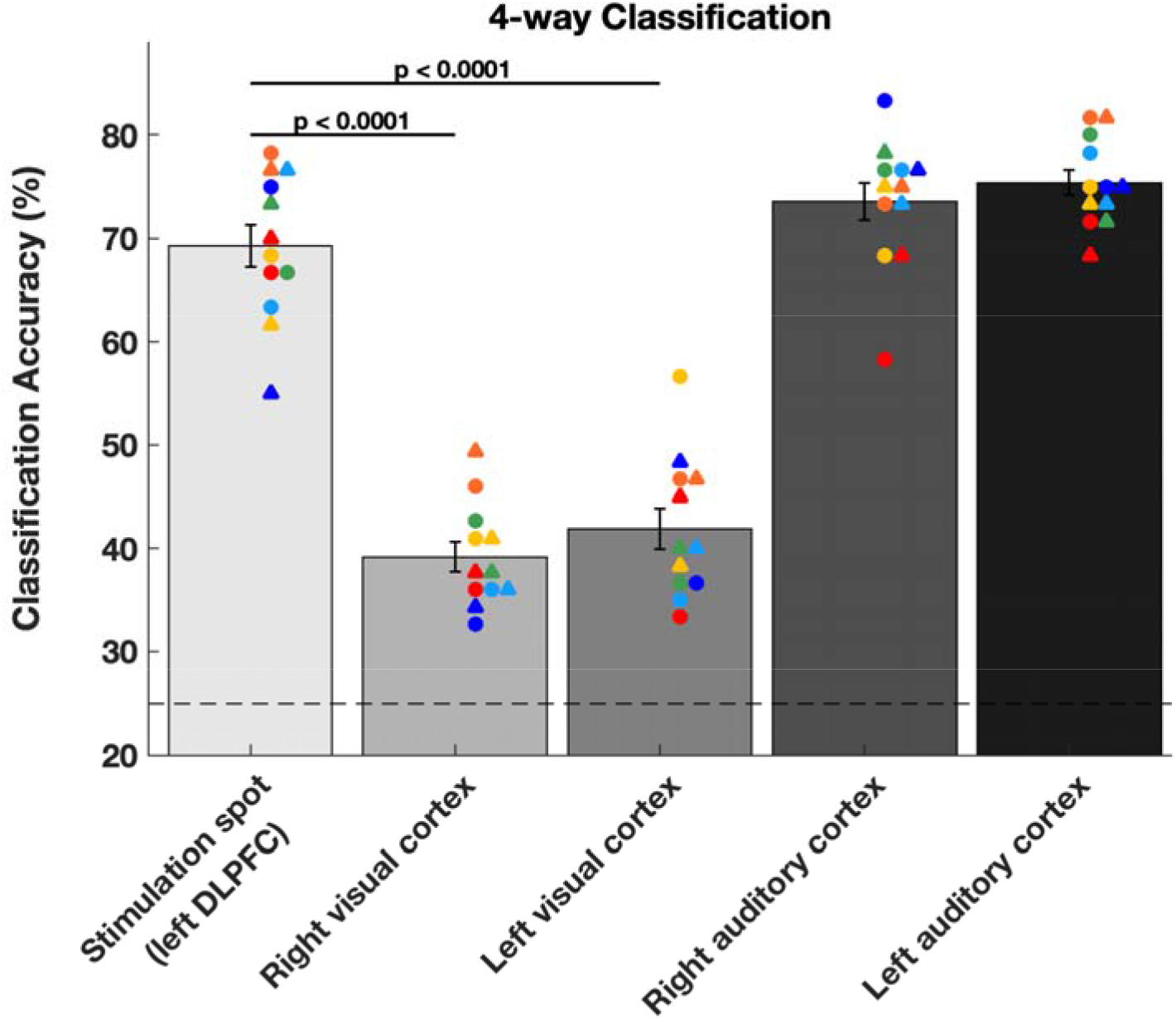
Decoding the TMS frequency using MVPA in different parts of the brain. We defined 20-mm spherical ROIs in left and right visual cortex, as well as left and right auditory cortex. The decoding performance at the stimulation spot was significantly greater than the decoding performance in the left and right visual cortex. In addition, decoding performance was very high in both left and right auditory cortex presumably due to the clicking sounds produced by the TMS pulses. Error bars represent SEM, colors represent unique subjects with data from Day 2 plotted as a circle and data from Day 3 plotted as a diamond. Chance performance is 25% (dashed line).

